# Association of a biomarker-based frailty index with telomere length in older US adults: Findings from NHANES 1999-2002

**DOI:** 10.1101/191023

**Authors:** Ghalib A. Bello, Yueh-Hsiu M. Chiu, Gerard G. Dumancas

## Abstract

**Objectives:** To study the link between frailty and cellular senescence, we examine the association of leukocyte telomere length (LTL) with a recently introduced measure of subclinical frailty that is based entirely on laboratory test biomarkers (FI-LAB).

**Methods:** This study was conducted on a random sample of 1890 Americans aged 60+. Multiple Linear Regression was used to examine the relationship between FI-LAB and LTL.

**Results:** A statistically significant association was found between FI-LAB and LTL after adjusting for multiple covariates, indicating that higher FI-LAB scores are associated with shorter telomeres.

**Discussion:** Our study results establish a link between subclinical frailty (FI-LAB) and cellular aging, which may help elucidate the pathophysiological mechanisms giving rise to frailty.

## INTRODUCTION

The physiological mechanisms underlying the link between telomere shortening and aging-related health conditions have not been completely elucidated. To this end, studies have attempted to examine the association between telomere length and aging-related parameters. One particular parameter that has received considerable attention is frailty, a clinical construct characterizing and quantifying the cumulative burden of aging-related health deficits. The 2 common measures of frailty (the “Frailty Phenotype” (FP) and “Frailty Index” (FI)) are typically constructed by tallying clinical symptoms of aging (e.g. diseases, disabilities, functional impairments), with higher scores indicating higher frequency and accumulation of age-related adverse health conditions (Xue, 2011). So far, studies examining the relationship of these clinical frailty measures with telomere length have found no association (Breitling et al., 2016; Collerton et al., 2012; Marzetti et al., 2014; Saum et al., 2014; Woo, Tang, Suen, Leung, & Leung, 2008; Yu, Tang, Leung, & Woo, 2015). This leaves open the question of how cellular senescence (of which telomere length is an indicator) eventually leads to clinical deficits seen among frail individuals.

Recently, Howlett et al (2014) introduced a new frailty metric that is derived solely from laboratory test and blood pressure abnormalities (Howlett, Rockwood, Mitnitski, & Rockwood, 2014). This index (named “FI-LAB” but abbreviated henceforth as FL) is constructed by computing the proportion of laboratory test biomarkers/physiological parameters on which an individual falls outside of the normal reference range. Studies have shown that while FL exhibits good agreement with clinical FIs, it appears to provide independent/complementary information about frailty, e.g. it predicts mortality independently of clinical FI (Howlett et al., 2014; Mitnitski et al., 2015; Rockwood, McMillan, Mitnitski, & Howlett, 2015). FL represents a novel and potentially useful approach to quantifying frailty, in that it focuses on ‘sub-clinical’ deficits (biological/physiological dysregulation), rather than clinically-detectable deficits (disabilities, functional impairments, etc.). The sub-clinical dysregulation (e.g. organ-level dysfunction) that FL measures is recognized as an intermediate stage in the process by which cellular-/molecular-level damage (evidenced by telomere attrition, etc.) scales up to and eventually culminates in clinically-visible health deficits (Blodgett, Theou, Howlett, Wu, & Rockwood, 2016; Zaslavsky, Cochrane, Thompson, Woods, & LaCroix, 2012). Examining the relationship of FL with telomere length could shed light on this process, as it would elucidate the intermediary role of sub-clinical dysregulation. In the current study, we assess the association of the novel frailty measure FL with leukocyte telomere length (LTL) among a randomly selected sample of the U.S. general population aged ≥60 years.

## METHODS

### Study Sample

The National Health and Nutrition Examination Survey (NHANES) is a program of population studies designed to evaluate the health and nutritional status of children and adults in the United States. This annual survey uses a stratified, multistage sampling approach to draw a random, representative sample of the non-institutionalized civilian US population (from all 50 states and the District of Columbia) (CDC (Centers for Disease Control and Protection), 2002). In this study, we use data collected in the 1999-2000 and 2001-2002 NHANES cycles. Over these 2 cycles there were 3706 participants aged ≥60 years. Informed consent was obtained from all NHANES participants and the Institutional Review Board of the NCHS approved the protocol.

### Computing FI-LAB (FL)

FL is computed as the proportion of biomarkers/physiological parameters on which an individual falls outside of the normal/clinical reference range, as proposed in a recent study by Howlett et al (2014) (Howlett et al., 2014). Only 21 out of 23 biomarkers/physiological parameters used in the Howlett et al (2014) study are available in NHANES 1999-2002 database. The 2 missing biomarkers were Free Thyroxine (T4) and syphilis antibody levels. Therefore the present study used 21 biomarkers to calculate each individual's FL score (Table 1).

**Table 1:** Computation of Frailty-Labs (FL)*

| Parameter | Normal/Reference Range |
| --- | --- |
| Albumin (g/dL) | 3.2 – 4.5 |
| AST/SGOT (U/L) | 8 – 33 |
| Systolic BP (mmHg) | 90 – 140 |
| Diastolic BP (mmHg) | 60 – 90 |
| Calcium (mg/dL) | 9.0 – 10.85 |
| Creatinine (mg/dL) | 0.6 – 1.2 |
| Glycohemoglobin/A1c (%) | < 6.5 |
| Hemoglobin (g/dL) | 13.5 – 18 |
| Mean Corpuscular Volume (fL) | 80 – 96 |
| Alkaline Phosphatase (U/L) | 20 – 130 |
| Phosphate (mg/dL) | 2.3 – 4.71 |
| Potassium (mEq/L) | 3.8 – 5 |
| Protein [total] (g/dL) | 6 – 7.8 |
| Sodium (mEq/L) | 136 – 142 |
| Urea Nitrogen [serum] (mg/dL) | 8 – 23 |
| White Blood Cell count (SI) | 1.8 – 7.8 |
| Folate (nM) | 11 – 57 |
| Folate, RBC (nM) | 376 – 1450 |
| TSH (μU/L) | 0.5 – 5 |
| Thyroxine [T4] (nM) | 71 – 161 |
| Vitamin B12 (pg/L) | 118 – 701 |

### Leukocyte Telomere Length

Telomere length assay was performed for NHANES 1999-2002 participants aged 20 years and older who had blood collected for DNA purification. Quantitative Polymerase Chain Reaction (PCR) was used to measure telomere length. To reduce measurement error, the following procedure was used - each sample was assayed 3 times on 3 different days, and telomere length measurements were standardized by dividing by standard reference DNA values (T/S ratio). Then the mean and standard deviation of the T/S ratios (across the 3 measurements) were computed for each sample (full details of this procedure are described elsewhere (Cawthon, 2002)).

### Covariates

Our analyses (described in the next section) utilized age, sex, race/ethnicity, socioeconomic status, education and tobacco exposure. Age (in years) at the time of NHANES participation was determined via a screening interview. To avoid disclosure risk, the ages of participants 85 years or older were reported as ‘85’. Sex, race and educational attainment were determined from survey questionnaire responses. Poverty income ratio was used as an indicator of socioeconomic status. It represents the ratio of annual family income to the poverty threshold. Poverty threshold income was derived from the guidelines outlined by the Department of Health and Human Services, and varied based on family size (US Department of Health & Human Services, 2011). Computed values of poverty income ratio that were above 5 (i.e. annual family income exceeding 5 times the poverty threshold) were recoded as ‘5’ in NHANES to avoid disclosure risk (CDC (Centers for Disease Control and Protection), 2002). The final poverty income ratio values ranged from 0 (lowest income) to 5 (highest income bracket). Serum cotinine concentrations (used here to quantify the extent of tobacco exposure) were measured in NHANES participants using isotope dilution high performance liquid chromatography/atmospheric pressure chemical ionization tandem mass spectrometry (ID HPLC-APCI MS/MS) (CDC (Centers for Disease Control and Protection), 2002).

### Statistical Analysis

To examine the association of FL with LTL, we used Ordinary Least Squares (OLS) regression. LTL (expressed as mean T/S ratio) was (natural) log-transformed and treated as the dependent variable in each model, with FL as a continuous explanatory/independent variable. This model was adjusted for age, sex, race/ethnicity (*White, Black, Hispanic* and *Other*), education (*less than high school education, and high school or more*), poverty income ratio (ratio of annual income to poverty threshold, treated as continuous), and serum cotinine levels (natural log-transformed).

The statistical analysis carried out in R statistical software (version 3.4.0, Vienna, Austria) (R Core Team, 2015).

## RESULTS

In the 1999-2000 and 2001-2002 cohorts, there were a total of 2667 subjects aged ≥60 years who had non-missing values of telomere length. Of these subjects, 777 were excluded due to missing values on other variables used in this study. This left a sample of n=1890. Table 2 provides summary statistics for demographic characteristics of this sample. The mean age of our analytic sample was 70.8 (SD: 7.7 years). The sample was 48% female. The breakdown of race/ethnicity was 62% white, 14% black and 22% Hispanic. 61% of members in our sample had a high school degree or higher, and the median ratio of annual income to the poverty threshold income was 2.4.

**Table 2:**
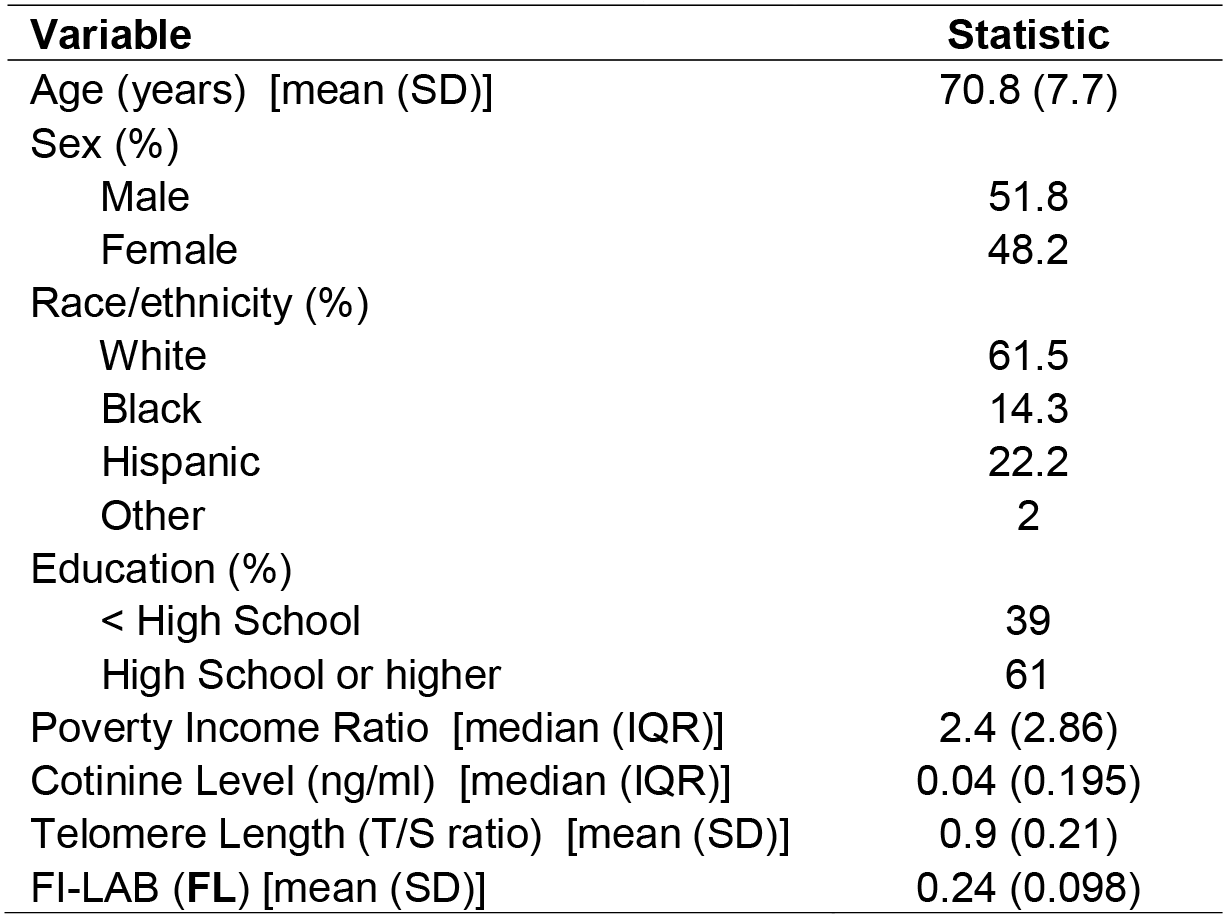
Characteristics of analytic sample from NHANES 1999-2002 (n=1890)

For FL, the mean score was 0.24, with a range of 0.05-0.63. Figure 1 shows histograms and bar plots for FL and LTL levels (expressed as mean T/S ratio) in the analytic sample. Mean T/S ratio had an average value of 0.9 in the analytic sample.

**Figure 1:**
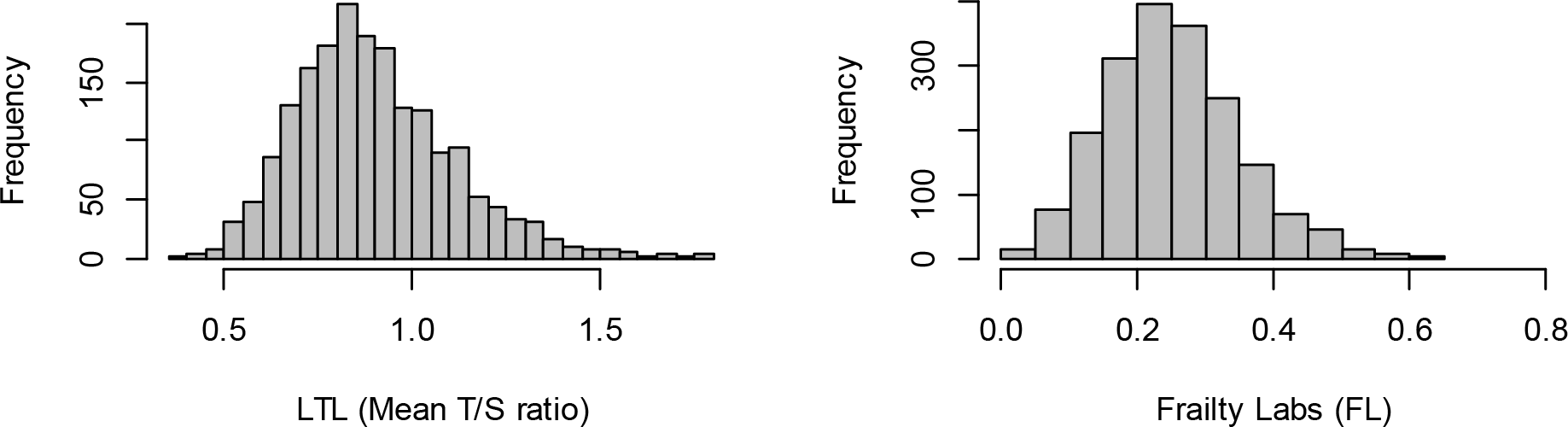
Sample distributions of LTL and FL

Table 3 provides results of the regression examining the association of LTL with FL, adjusting for covariates. Coefficient estimates (and SEs) and p-values are provided. After adjusting for age, sex, race/ethnicity, education, poverty income ratio and cotinine levels, FL demonstrated a significant association with telomere length (p = 0.0056). As expected, age had a negative and significant association with LTL (p<.0001). Female sex was significantly positively associated with higher telomere length (p<.0001), which is in line with multiple studies that have shown that females, on average, have longer telomeres than males (Gardner et al., 2014). Poverty income ratio showed a significant positive association with LTL (p=.011), suggesting that, after controlling for other potential confounders, higher income is associated with longer LTL.

**Table 3:** Linear Regression examining association between LTL (expressed as Mean T/S ratio) and FL, adjusting for demographic and lifestyle-related covariates

| Variable |  | Beta | SE | p-value |
| --- | --- | --- | --- | --- |
| Intercept |  | 0.317 | 0.0565 | <.0001 |
| FL |  | -0.146 | 0.0526 | 0.0056 |
| Age |  | -0.007 | 0.0007 | <.0001 |
| Sex | *Male* | *Ref* |  |  |
|  | *Female* | 0.053 | 0.0103 | <.0001 |
| Race | *White* | *Ref* |  |  |
|  | *Black* | 0.022 | 0.0157 | 0.171 |
|  | *Hispanic* | -0.012 | 0.0144 | 0.395 |
|  | *Other* | -0.019 | 0.0369 | 0.608 |
| Poverty income ratio |  | 0.010 | 0.0039 | 0.011 |
| Education | *Less than HS* | *Ref* |  |  |
|  | *HS or higher* | 0.006 | 0.0124 | 0.651 |
| log(cotinine) |  | -0.003 | 0.0526 | 0.092 |

\*Beta: Maximum Likelihood Estimate (MLE) of coefficient corresponding to frailty variable
\*SE: Standard Error of beta coefficient estimate

## DISCUSSION

Studies have shown both telomere length and frailty are determinants of aging-related health endpoints. However, a definitive association between these two correlates of aging has not been established - multiple studies investigating this link found no association. In this study, we reexamined this question using a novel laboratory measure-based index of frailty (FL) which focuses on sub-clinical/pre-clinical deficits, in contrast to previously commonly used frailty measures (FI and FP) which are largely based on clinically observable deficits, and which have not demonstrated significant associations with telomere length (Breitling et al., 2016; Collerton et al., 2012; Marzetti et al., 2014; Saum et al., 2014; Woo et al., 2008; Yu et al., 2015). In a covariate-adjusted model, we found a significant association between increased FL and decreased LTL.

These results can be interpreted in the context of a conceptual model that views frailty as a multilevel cascade of pathophysiological processes that begins at the cellular level (oxidative damage, telomere attrition), progresses to the system/organ level (e.g. systemic inflammation, metabolic dysfunction) and culminates in clinically evident health endpoints (like diseases and functional impairments) (Zaslavsky et al., 2012). However, prior studies have failed to find any association between telomere length and these clinical endpoints (as quantified by FI and FP). Our finding that FL (sub-clinical frailty) is associated with telomere length confirms that the systemic/organ-level physiological dysregulation that precedes clinical endpoints has a more direct relationship with cellular senescence.

It is worth nothing that FL bears close similarity to some of the biomarker-based measures of biological aging that have been developed in recent years (Bello & Dumancas, 2017; Belsky et al., 2015; Cohen, Milot, & Morissette-Thomas, 2013; Klemera & Doubal, 2006; Levine, 2013; Milot et al., 2014; Putin et al., 2016). These measures quantify biological aging by combining age-correlated clinical biomarkers (routinely measured in standard laboratory tests) into a unidimensional index correlated with chronological age (Levine, 2013). The aggregation of these biomarker measures into a composite index is done using regression-based or machine learning techniques that ‘weight’ each biomarker in relation to its strength of association with chronological age. These biological age measures have been shown to be correlated with mortality, adverse health outcomes, and telomere length (Kulminski, Ukraintseva, et al., 2008). The resemblance of FL to these biological age indices is evident, and it could in fact be argued that FL is a crude measure of biological age that dichotomizes its component biomarkers (based on clinical reference ranges) and assigns them equal weights. Like other measures of biological age, FL also exhibits an association with telomere length.

We highlight one further point that may shed more light on why FL is associated with telomere length but FI and FP are not. In multiple studies utilizing FI and FP, both measures indicate frailty is more prevalent in females than males (Gordon et al., 2017). This is at odds with the established consensus that telomeres tend to be longer in women (Gardner et al., 2014), and also that women have lower mortality risk compared to men of similar age. This observation that women are more frail than men but ultimately have lower mortality has been referred to as the 'male-female morbidity-mortality paradox’ (Kulminski, Culminskaya, et al., 2008), or the ‘male-female health-survival paradox’ (Gordon et al., 2017; Oksuzyan, Juel, Vaupel, & Christensen, 2008; Verburgge, 1985). However, compared to FI and FP, studies find that the FL shows women to be relatively less frail (Howlett et al., 2014). This makes FL more in agreement with telomere length than FI/FP, the clinical measures of frailty. Taken together, these observations suggest that while females appear to experience greater number of clinically detectable (macroscopic) functional deficits (compared to males), this trend appears to be diminished or reversed at the cellular/molecular and organ/systemic levels.

The main strength of this study is the large sample size and the representative nature of the cohort. NHANES participants are randomly selected from the general population of noninstitutionalized US civilians. We also acknowledge a few limitations. A major limitation is that NHANES is cross-sectional in nature, and in particular, LTL was measured at a single time-point, meaning that telomere attrition rate could not be examined. Single time-point measurements of telomere length may be limited in their ability to capture the cellular aging process (Aviv et al., 2009). Further, there may be residual confounding by socioeconomic status (in addition to education and income that were adjusted in our analysis) or unmeasured confounding by factors that may be associated with both FL and LTL, such as diet intake or exposure to environmental toxicants. Nonetheless, rather than these dietary or environmental factors being the confounders, it is more likely that FL may be on the pathway of the association between these factors and telomere length. Finally, out of the 23 biomarkers and physiological parameters used to construct FL (see (Howlett et al., 2014)), only 21 were available in the NHANES 1999-2002 data, thus our FL was constructed using these biomarkers. The 2 unavailable biomarkers were Free Thyroxine (T4) and the syphilis antibody titer from the VDRL (Venereal Disease Research Laboratory) screening test (Hart, 1986). Finally, our study was limited only to the U.S. population, therefore future studies in other geographic areas are warranted to replicate our findings of a significant association between FL and LTL.

While FL has not gained widespread adoption as a frailty measure, some of its practical benefits are worth highlighting here. Because it is largely based on standard laboratory tests, it is a viable alternative to commonly used operational frailty metrics which are based largely or solely on clinical deficits, many of which are determined via subjective self-report. Lab tests, on the other hand, are a standard and objective measure of health deficits which do not require self-assessment on the part of individuals. FL can be constructed from standard lab tests (and blood pressure measurements) which are readily available in many health/clinical databases and are often a routine part of standard care. Studies report that FL shows good agreement with other frailty measures (Ritt, Jager, Ritt, Sieber, & Gaßmann, 2017) and is strongly associated with mortality, hospital and healthcare utilization, medication use, and self-reported health (Blodgett et al., 2016; Howlett et al., 2014).

In conclusion, we have demonstrated that a frailty measure constructed from standard laboratory tests and physiological parameters exhibits a significant association with LTL among a cohort of community-dwelling members of the U.S. general population aged 60 and older. Our study highlights unique and useful features of the FL that set it apart from clinical measures of frailty, and could potentially help in elucidating the pathophysiological mechanisms of frailty.

## ACKNOWLEDGMENTS

None

